## Supplementary figures and images for "Evolutionary trade-off and mutational bias could favor transcriptional over translational divergence within paralog pairs"

### S1 File

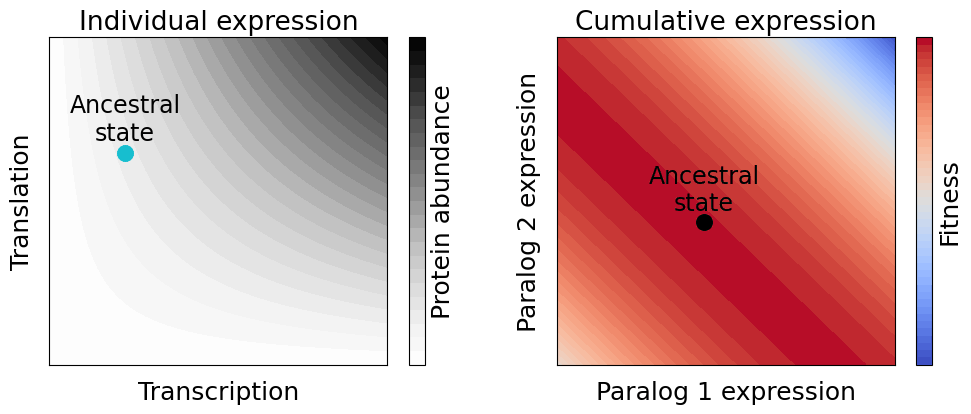
